# Different folding mechanisms in prion proteins from mammals with different disease susceptibility observed at the single-molecule level

**DOI:** 10.1101/2024.08.09.607387

**Authors:** Uttam Anand, Shubhadeep Patra, Rohith Vedhthaanth Sekar, Craig R. Garen, Michael T. Woodside

## Abstract

Misfolding of the protein PrP causes prion diseases in mammals. Disease susceptibility varies widely among species, despite PrP sequences differing by only a few amino acids. How these differences alter PrP folding and misfolding remains unclear. We compared the folding dynamics of single PrP molecules from three species with different disease susceptibility: dogs (immune), hamsters (susceptible), and bank voles (extremely susceptible). Measurements with optical tweezers revealed important differences between the folding cooperativity, pathways, energy barriers, and kinetics of these proteins. In contrast to the two-state folding of hamster PrP, dog PrP always folded through multiple intermediates. However, both featured rapid native folding, homogeneous energy barriers, and no readily observable misfolding. Bank vole PrP also folded via intermediates, but more slowly and via inhomogeneous barriers. Most notably, it formed several metastable misfolded states starting from the unfolded state. Analyzing the sequence of intermediates seen in pulling curves, we found significant differences in the folding pathways for dog and bank vole PrP, implying that sequence mutations altered energy barriers so as to redirect folding pathways. These results show that subtle differences in PrP sequence between species produce profound changes in folding behavior, providing insight into the factors underlying misfolding propensity.

---

Prion diseases are a family of fatal neurodegenerative disorders that affect a wide range of mammals, including Cretzfeldt-Jakob disease in humans, scrapie in sheep, bovine spongiform encephalopathy in cattle, and chronic wasting disease in cervids.^1,2^ The central pathogenic event in these diseases is the conversion of the membrane-associated prion protein from its native cellular form, denoted PrP^C^, into a β-rich misfolded isoform, denoted PrP^Sc^, that is neurotoxic.^3–6^ Crucially, PrP^Sc^ can recruit other PrP^C^ molecules to propagate the misfolding through a templated mechanism, acting as a form of infectious agent that can transmit disease.^1,7^ Similar templated propagation of misfolding is seen in other proteins associated with various neurodegenerative diseases, including Alzheimer’s, Parkinson’s, and amyotrophic lateral sclerosis, indicating that prion-like conversion is not limited to PrP.^8,9^ However, the mechanisms by which these proteins misfold and induce template propagation of misfolding remain elusive.^10,11^

Intriguingly, although PrP is highly conserved across the animal kingdom,^12–14^ with a high degree of sequence identity across mammals, different species show a wide range of variations in susceptibility to prion diseases. Some species appear to be immune or very resistant to propagating prions (*e*.*g*. dogs, horses^15–17^), some can support infection but only with difficulty (*e*.*g*. rabbits, pigs^18–22^), whereas others are much more susceptible (*e*.*g*. rodents, cattle, sheep, deer, cats, primates^22–25^). Studies using animals from resistant species modified transgenically to express PrP from susceptible species imply that resistance/susceptibility arises mainly from the PrP sequence,^26^ even though the natively folded structure of PrP is very similar across all these species.^27–30^ Previous work has explored various properties of PrP that might explain differential disease susceptibility. For example, specific residue mutations have been identified that contribute to susceptibility or resistance,^17,20,31,32^ differences in local structural flexibility within the native fold have been linked to differential susceptibility,^33–35^ and both the propensity of native PrP to reconfigure into β-rich isoforms^36^ and the reconfiguration dynamics of unfolded PrP^37^ correlate to some extent with susceptibility. These results suggest that the sequence differences perturb the folding energy landscape so as to enhance or suppress disease-relevant misfolding. However, the mechanisms driving differences in disease susceptibility remain unclear, in part because the folding dynamics, pathways, and energy landscapes have never been compared systematically for PrPs from species with different disease susceptibility.

Single-molecule methods offer an attractive approach for addressing this question.^38,39^ By directly monitoring the folding trajectories of individual molecules, they can detect and characterize transient intermediates and rare misfolded states,^40^ map folding/misfolding pathways,^41^ and distinguish and characterize different subpopulations of behavior.^42–45^ Single-molecule force spectroscopy (SMFS), in which force applied to a molecule and the resulting changes in structure are observed,^46^ is particularly useful for this purpose, as it allows the properties of the energy land-scape underlying the folding dynamics to be quantified and manipulated.^47^ SMFS has been used previously to probe folding/misfolding pathways and energy landscapes in monomers and dimers of hamster PrP,^48–50^ the formation and properties of dimers in mouse and human PrP,^51,52^ the properties of human PrP within amyloids,^53^ and the effects of anti-prion compounds on hamster PrP folding.^54,55^ This work has provided important insights into the folding of individual PrP molecules, but the behavior of PrP from different species has not yet been compared systematically under the same conditions.

We used SMFS to do so, extending previous work on hamster PrP^48,49^ to compare the folding of PrP from three species with different disease susceptibility: dog, hamster, and bank vole. Dogs appear to be immune from prion diseases;^56^ in contrast, bank voles are the most susceptible species known, and can be infected by all prions from any source.^57^ Hamsters have intermediate susceptibility: they were the first animal model used to study prion diseases in the lab, but they can only be infected by a limited set of prions.^58^ Despite these differences, all three PrP variants share the same fold in the structured C-terminal domain^28–30^ and differ by only a few amino acids in sequence (Fig. 1A). Using optical tweezers to unfold and refold single PrP molecules, we found that each variant featured quite distinct folding behavior: multi-state pathways for dog PrP (CaPrP) and bank vole PrP (BvPrP) in contrast to two-state folding for hamster PrP (HaPrP), folding pathways with different intermediates for CaPrP compared to BvPrP, homogeneous energy barriers for CaPrP and HaPrP but not BvPrP, and metastable misfolded states for BvPrP but not the others. These results show that changing only few amino acids in PrP can lead to dramatic changes in folding and misfolding without altering the final native structure.

**Fig. 1:**
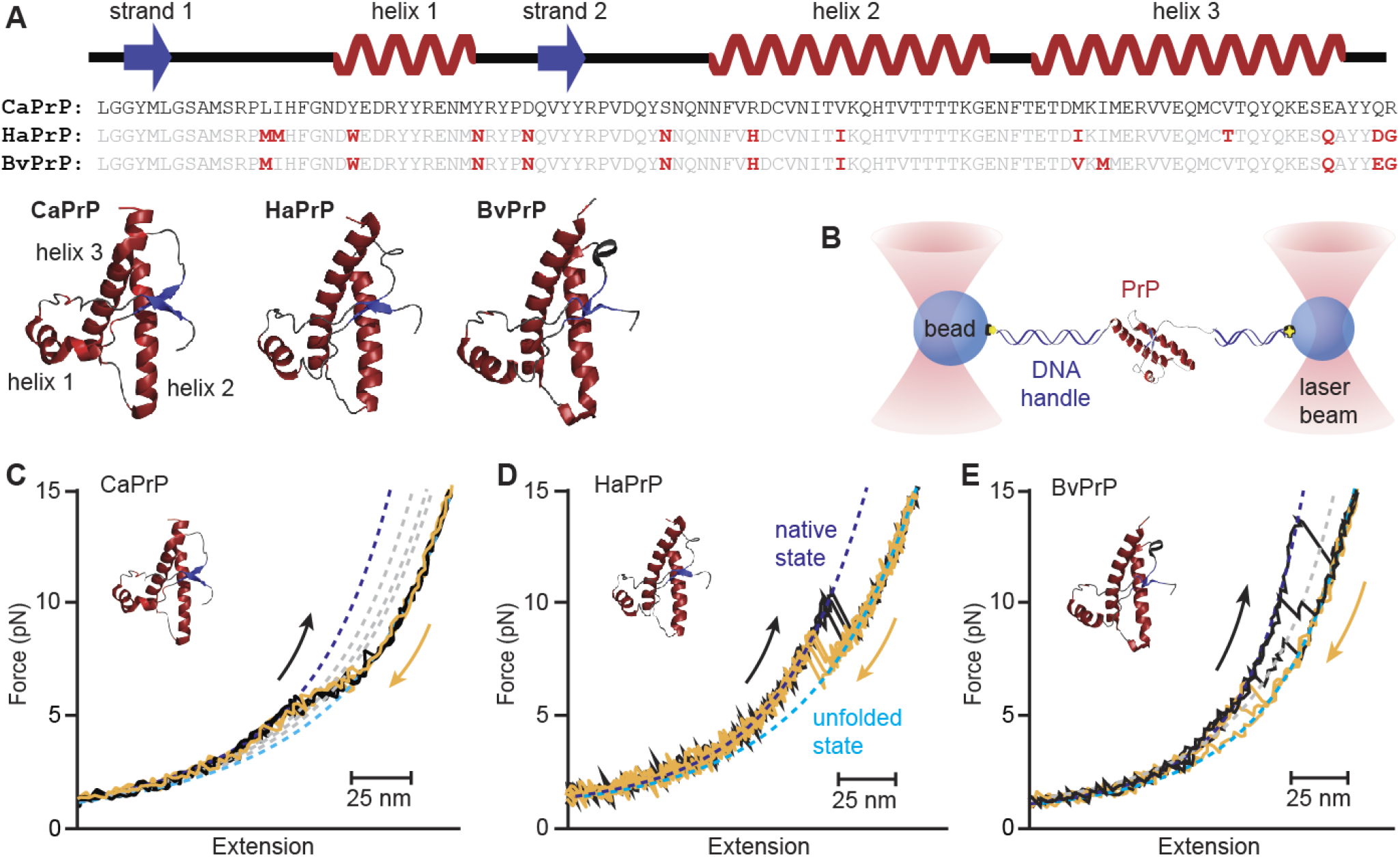
Force spectroscopy of PrP from species with different disease susceptibility. (A) Sequences, secondary structure, and NMR structures of the structured C-terminal domain of PrP from dogs (CaPrP), hamsters (HaPrP), and bank voles (BvPrP). Sequence differences from CaPrP shown in red. (B) Schematic of SMFS assay: a PrP molecule bound to DNA handles at its termini is attached to beads held in optical traps. (C)–(E) Representative force-extension curves (FECs) for (C) CaPrP, (D) HaPrP, and (E) BvPrP, showing unfolding (black) and refolding (gold) curves. Dashed lines: WLC fits to the folded (blue), intermediate (gray) and unfolded (cyan) states. (C) CaPrP folds/unfolds in quasi-equilibrium over a narrow force range via multiple intermediate states. (D) HaPrP folds/unfolds over a narrow force range in a two-state process slightly out of equilibrium. (E) BvPrP folds/unfolds out of equilibrium over a wide force range via at least one intermediate.

## RESULTS

### Native folding of PrP from different species

To measure the folding dynamics of single PrP molecules, we tethered recombinant PrP molecules attached covalently at their termini to double-stranded DNA handles between polystyrene beads held in dualbeam optical traps (Fig. 1B). We moved the traps apart at a constant speed to ramp up the force and unfold the protein, brought the traps together again to ramp down the force and refold the protein, and repeated the process to generate multiple cycles of successive force-extension curves (FECs), waiting 2–15 s at low force between cycles to allow time for the protein to refold. Representative unfolding and refolding FECs (Fig. 1C–E, respectively black and gold) show typical behavior: a non-monotonic rise of force with extension as the DNA handles are stretched out, interrupted by one or more ‘rips’ in which the extension jumps abruptly as part or all of the protein unfolds/refolds.

Each of these sets of FECs shows qualitatively distinct behavior. CaPrP (Fig. 1C) unfolded and refolded in a force range of ∼5–10 pN through a series of many small steps, indicating multiple intermediate states; unfolding and refolding curves lie on top of each other, implying that unfolding and refolding were fast and in quasi-equilibrium. HaPrP (Fig. 1D), as reported previously, unfolded and refolded in the range ∼6–12 pN without any intermediates but with some hysteresis, implying that it was out of equilibrium. BvPrP (Fig. 1E) unfolded and refolded typically through one intermediate, over a wider range of forces (∼5–20 pN), showing significant hysteresis. Notably, folding appeared to be significantly slower for BvPrP than the other two: whereas CaPrP and HaPrP always refolded into the initial state in refolding curves while still under tension, BvPrP did not always do so; longer waiting times between successive pulls increased the fraction of FECs returning to the initial state (Fig. S1).

The total contour-length change (Δ*L*_c_) upon unfolding/refolding in FECs reveals whether the folded structure is consistent with expectations for native folding. We found the total Δ*L*_c_ by fitting each branch of an FEC to an extensible worm-like chain (WLC) polymer elasticity model,^59^ described by Eq. 1 in Methods (Fig. 1C–E, dashed lines), and then measuring the change between most-folded and most-unfolded branches. The result was in each case consistent with expectations for unfolding/refolding of the native structure: 34.1 nm for CaPrP, 34.3 nm for HaPrP, and 34.6 nm for BvPrP, based on high-resolution NMR structures.^28–30^ For CaPrP and HaPrP, all FECs showed behavior similar to that illustrated in Fig. 1, with the total Δ*L*_c_ averaged over all FECs from all molecules found to be 34.9 ± 0.3 nm for CaPrP and 34.1 ± 0.4 nm for HaPrP (as reported previously^48^); errors represent standard error of the mean (Table S1). For BvPrP, in contrast, although the majority of FECs exhibited total Δ*L*_c_ consistent with native folding, with an average value of 34.5 ± 0.3 nm, ∼27% showed smaller values, implying that the protein did not refold into the native structure.

Given the profound differences in folding cooperativity between the three PrP variants, we studied the intermediate states for native unfolding and refolding in more detail. HaPrP, which showed cooperative twostate folding without evidence of intermediates,^48^ was omitted from this analysis. For both CaPrP and BvPrP, almost all (∼93%) of unfolding FECs contained at least one intermediate: most commonly two intermediates for CaPrP (Fig. 2A, top inset) but only one for BvPrP (Fig. 2B, top inset). Neither ever showed more than 3 intermediates. Refolding featured more FECs without intermediates (∼25% in each case), but most still showed 1–3 intermediates (Fig. 2, bottom insets). However, examining the Δ*L*_c_ values for intermediates observed in different FECs from a given protein, we found that the same intermediate states were not visited in every FEC: instead, each time the proteins unfolded or refolded, the intermediates populated were selected from among a larger set of possible states.

**Fig. 2:**
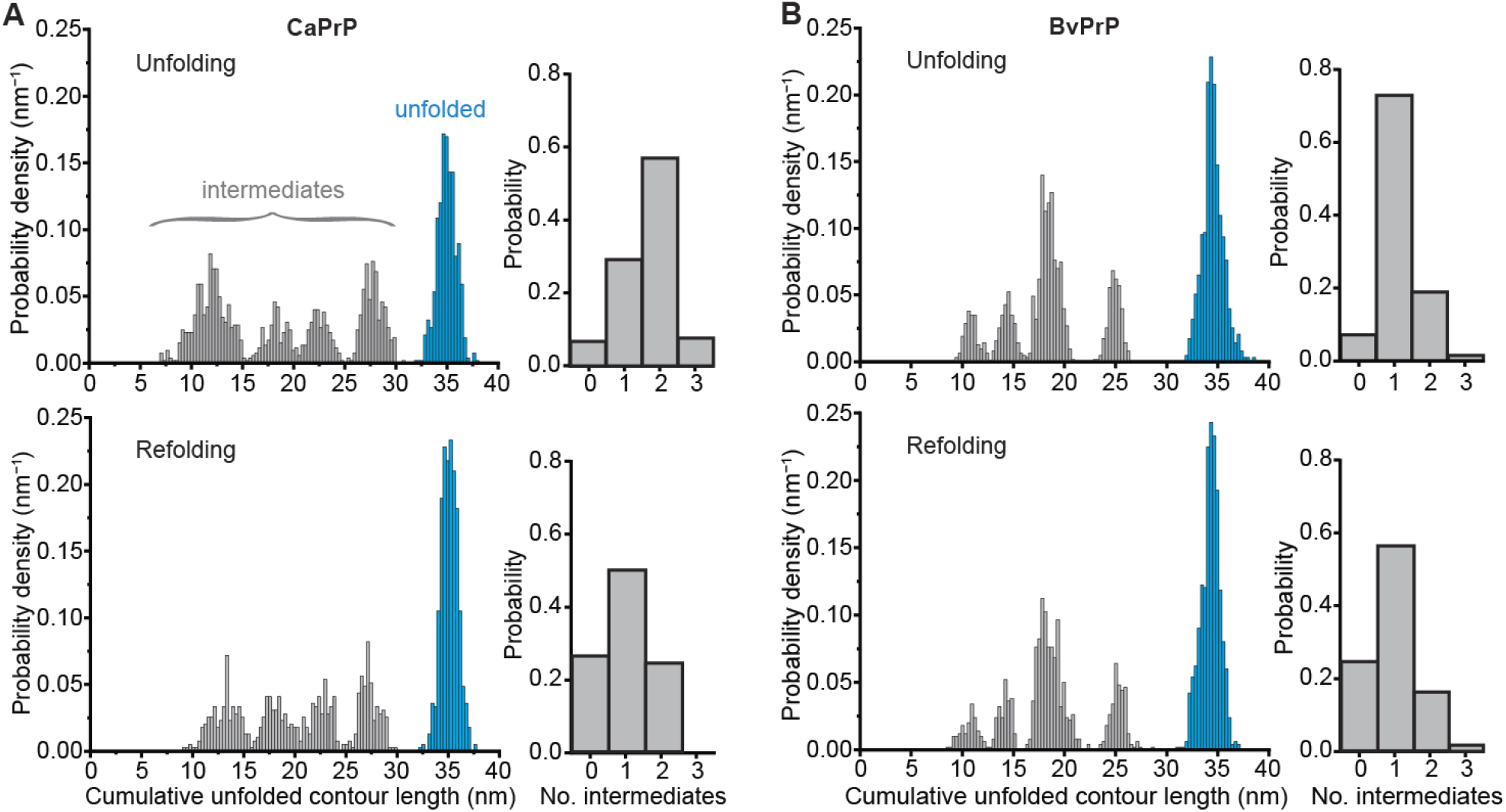
Intermediate states in CaPrP and BvPrP folding and unfolding. (A) Distribution of the unfolded contour length measured from WLC to all branches in all FECs for unfolding (top) and refolding (bottom) of CaPrP. Four peaks with different lengths representing at least four distinct intermediates are seen (grey), in addition to the peak for the unfolded state (cyan). Insets: distribution of number of intermediates seen in a given FEC for unfolding (top) and refolding (bottom). (B) Same for BvPrP. The lengths of the intermediate states and the number of intermediates differ from CaPrP.

Plotting the distribution of values for the contour length of unfolded protein (reflecting how much of the protein remained folded) seen in every branch of every FEC, as measured by WLC fits (Fig. 1C–E, dashed lines), we found four peaks for both CaPrP (Fig. 2A, grey) and BvPrP (Fig. 2B, grey), aside from the unfolded state peak (Fig. 2, cyan), indicating the existence of at least four structurally distinct intermediates for both proteins. The same peaks were seen in unfolding (Fig. 2, top) and refolding (Fig. 2, bottom) FECs. However, the positions of the peaks differed for CaPrP as compared to BvPrP, indicating that the intermediate structures formed were different. For CaPrP, the four peaks were denoted as intermediates I^Ca^_1_ through I^Ca^_4_, whereas for BvPrP they were denoted as intermediates I^Bv^_1_ through I^Bv^_4_. Unfolded contour lengths for all states of all proteins are listed in Table S1.

### Force distributions and energy landscape properties

The forces measured in SMFS contain information about the energy landscape underlying the folding behavior, such as the distance to the energy barrier, Δ*x*^‡^, and the barrier height, Δ*G*^‡^, in addition to information about the kinetics.^47^ To characterize energy landscape properties, we fitted the distribution of forces for the first unfolding rip (which reflects the first event disrupting the native structure) to a model that assumes unfolding from a homogeneous initial state.^60^ This model (Eq. 2 in Methods) was previously found to fit well to the unfolding force distribution for HaPrP.^49^ The unfolding force distribution for CaPrP was described well by this model (Fig. 3A), even though the unfolding forces were generally lower than for HaPrP (Fig. 3B) and represented unfolding of only part of the protein. The distance to the barrier was the same within error for both proteins: Δ*x*^‡^ = 10 ± 1 nm for CaPrP compared to 9 ± 1 nm for HaPrP; the barrier heights were also similar within error, at ΔG^‡^ = 77 ± 5 kJ/mol for CaPrP and 64 ± 6 kJ/mol for HaPrP. The most significant difference from the fits was the unfolding rate at zero force: *k*_unfold_ = 10^−3±1^ s^−1^ for CaPrP, much faster than for HaPrP with 10^−6±1^ s^−1^.

**Fig. 3:**
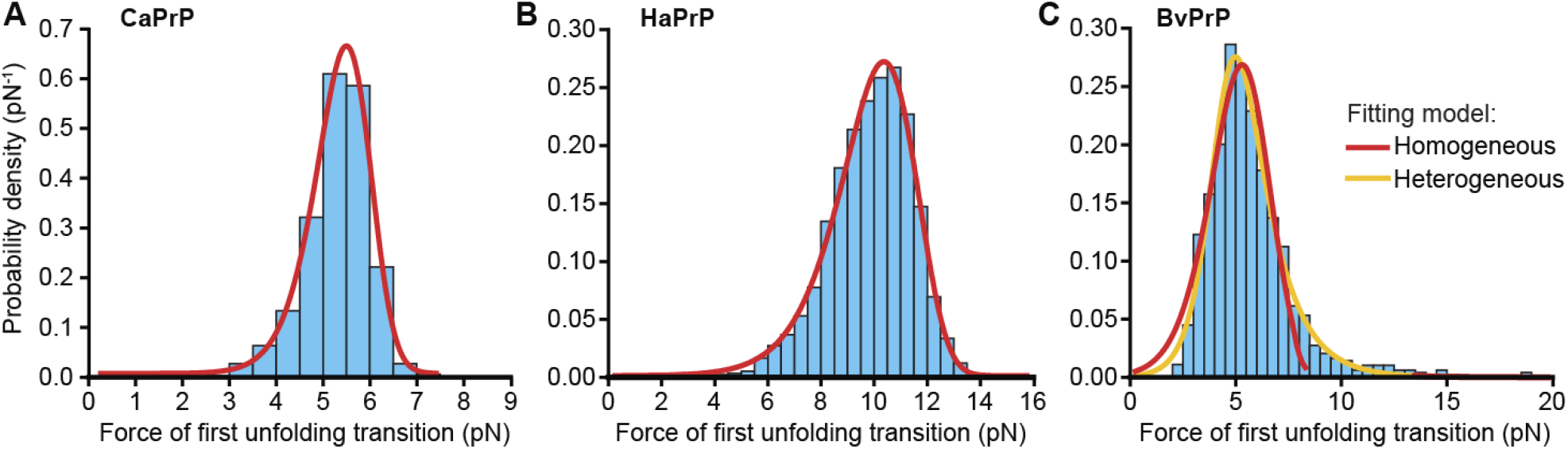
Unfolding force distributions. (A) Distribution of forces for the first unfolding transition in FECs of CaPrP. Red: fit to Eq. 2 (homogeneous barrier model). (B) Same for HaPrP. (C) Same for BvPrP. Red: fit to Eq. 2 (homogeneous barrier model), orange: fit to Eq. 3 (heterogeneous barrier model).

Turning to BvPrP, however, the distribution of forces for unfolding the state with native contour length (Fig. 3C) was not well fit by the same model (Fig. 3C, red): there was a long tail at high forces that was inconsistent with the distribution expected for a single initial state unfolding through a homogeneous barrier. Such a tail is instead characteristic of unfolding from heterogeneous states that can interconvert during unfolding and thus sample different barriers.^61^ Fitting instead to a model for heterogeneous unfolding (Eq. 3), which parametrizes the distribution by Δ*x*^‡^, *k*_unfold_, and a heterogeneity parameter *Δ*,^61^ we found excellent agreement (Fig. 3C, orange), with Δ*x*^‡^ = 5.5 ± 0.3 nm, *k*_unfold_ = 10^−2±1^ s^−1^, and *Δ* = 1.7 ± 0.2. The considerably smaller value of Δ*x*^‡^ compared to HaPrP and CaPrP indicates that BvPrP is more rigid mechanically, despite the similar overall structure of the folded state. Because the heterogeneous model does not return a barrier height, no comparison can be made for Δ*G*^‡^, but it must be comparatively high given the significant hysteresis in the pulling curves, and the nature of the barrier is presumably qualitatively different given that BvPrP samples heterogeneous barriers rather than a single well-defined barrier as in CaPrP and HaPrP.

### Native folding pathways

We next analyzed the sequence of intermediates observed in FECs to deduce the pathways for native folding/unfolding. For each FEC, we assigned every branch of the curve to either the native state (N), the unfolded state (U), or an intermediate state (one of the 4 candidates each for CaPrP and BvPrP; not an option for HaPrP), based on the contour lengths from WLC fits. We then counted up all the instances of transitions between any pair of states to create transition maps for unfolding and refolding (Fig. 4A–C). The transition maps reveal all pathways that the protein followed through the available intermediates when changing between folded and unfolded states, as well as the statistical weights for those pathways. For HaPrP (Fig. 4B), the transition map is trivial as there are no intermediates. For CaPrP and BvPrP, however, the maps are complex, featuring multiple possible pathways through the intermediate states. Crucially, the maps are very different for CaPrP and BvPrP. There is no single dominant pathway for CaPrP (Fig. 4A), although unfolding proceeds most commonly through the intermediates I^Ca^_1_ and I^Ca^_4_. In contrast, for BvPrP (Fig. 4C), the pathway F ↔ I^Bv^_3_ ↔ U clearly dominates the transition map, revealing the intermediate I^Bv^_3_ as crucial for both native unfolding and refolding.

**Fig. 4:**
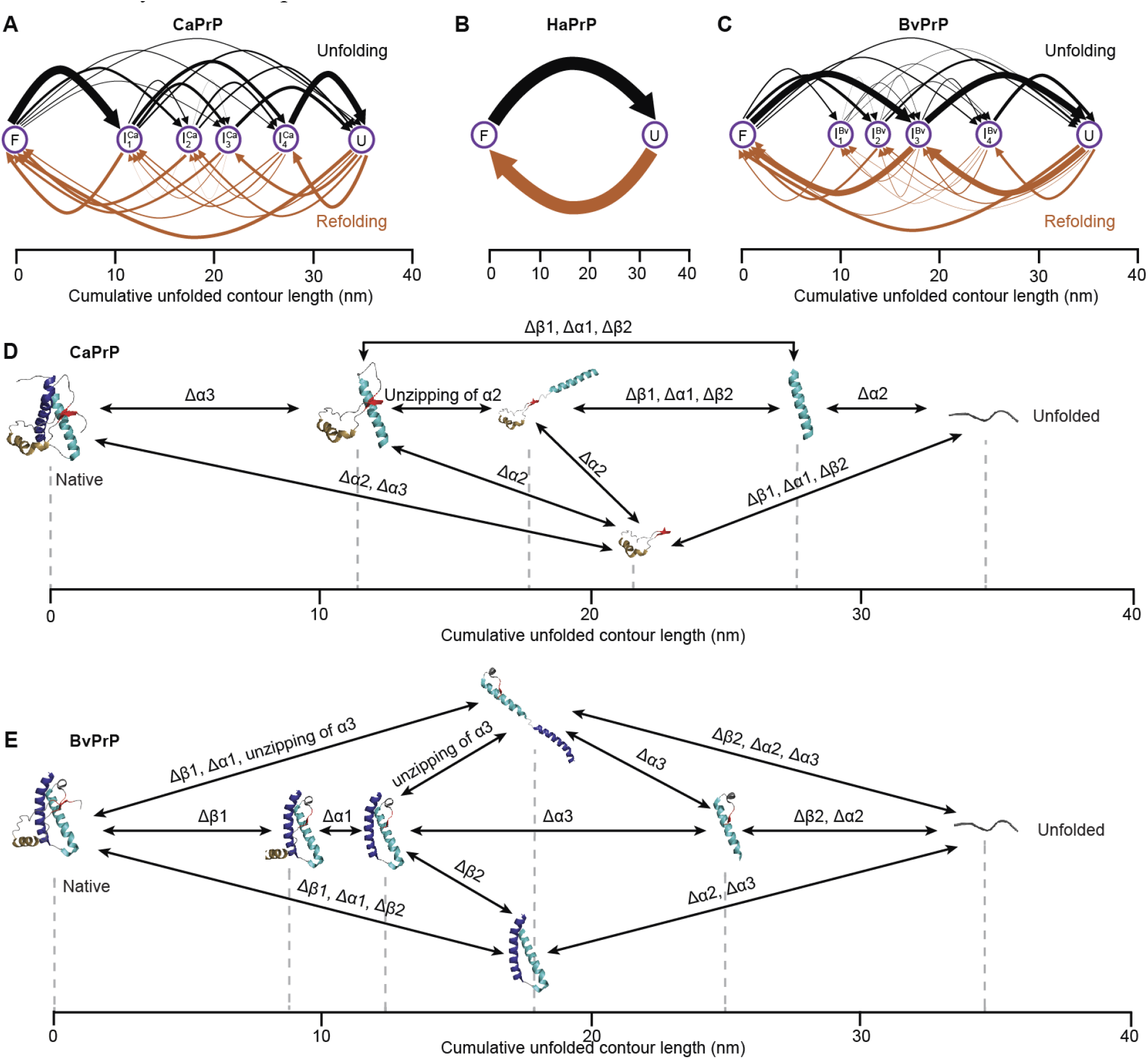
Native folding pathways. (A–C) Transition maps for unfolding (black) and refolding (gold) of (A) CaPrP, (B) HaPrP, and (C) BvPrP. Arrow thickness proportional to relative frequency of transition. (D–E) Native folding pathways showing possible intermediate state structures for (D) CaPrP and (E) BvPrP, as deduced from network analysis of intermediate-state contour lengths. Dashed lines show contour lengths of observed states.

To interpret the transition maps structurally, we explored what substructures might be associated with the intermediate states by relating the length changes seen in FECs with the lengths predicted from unfolding different structural elements of the native conformation, a method applied previously in SMFS studies of other proteins.^40,41,62^ We first catalogued all plausible partially native intermediate states (*i*.*e*., intermediates retaining some fraction of the secondary structure elements of the native state). To do so, we assumed that all the loops in PrP are unstable, each of the three α-helices are stable by themselves or in combination, the two β-strands are stable together but not individually, and strand 2 is stable in contact with helix 2. We verified these assumptions with all-atom MD simulations, finding that these substructures persisted in simulation up to 750 ns long (Fig. S2). The expected unfolded length for every structure is listed in Tables S2. For each FEC, we then compared the observed *L*_c_ values to the lengths for each substructure, identifying the set of possible structures that could explain the observed *L*_c_ values. Based on the sequence of transitions in each FEC, we identified possible sequences of sub-structures (*i*.*e*., folding/unfolding pathways) consistent with that FEC, under the constraint that once a particular element of the secondary structure had unfolded it could not later refold. We repeated this analysis for all FECs to find the minimal set of structures needed to account for the major fraction of FECs. The resulting pathways, shown in Figs. 4D–E, are able to account for ∼ 81% of FECs for CaPrP and ∼85% for BvPrP; adding more pathways did not substantially increase the number of FECs consistent with the model (Fig. S3). Intriguingly, this analysis suggests that the most common initial step in CaPrP folding (state I^Ca^_4_) involves forming a single large helix (Fig. 4D), in contrast to the dominant folding pathway for BvPrP (passing through I^Bv^_3_), which involves first forming both helices 2 and 3, or else helices 1 and 2 plus strand 2 (Fig. 4E). Another notable difference suggested by the analysis is that the β-sheet/helix 1 complex forms only in the last step for BvPrP, instead of much earlier for CaPrP.

### Multiple metastable misfolded states for bank vole PrP

Whereas all CaPrP and HaPrP FECs refolded into the native state, not all did so for BvPrP: in 27% of them, the total Δ*L*_c_ upon unfolding was shorter than native (Fig. 5A, red), indicating unfolding from a state other than the native fold. Such non-native Δ*L*_c_ could indicate a molecule still stuck in an intermediate on the native folding pathway, owing to the slow folding kinetics of BvPrP, or alternatively a molecule trapped in a misfolded state (a metastable state prevented from proceeding to the native state by large energy barriers^63^). Evidence for slow refolding of BvPrP was seen in refolding-unfolding cycles, where refolding FECs sometimes ended in a partially folded state consistent with an intermediate on the native folding pathway, and the subsequent unfolding FECs either showed unfolding from the same state (native fold still incomplete) or else unfolding with native total Δ*L*_c_ (native folding completed during the waiting time between pulls). Evidence for misfolding would instead consist of FECs where the protein was trapped in a state with Δ*L*_c_ shorter than the native length but inconsistent with any of the intermediate states on the native pathway.

**Fig. 5:**
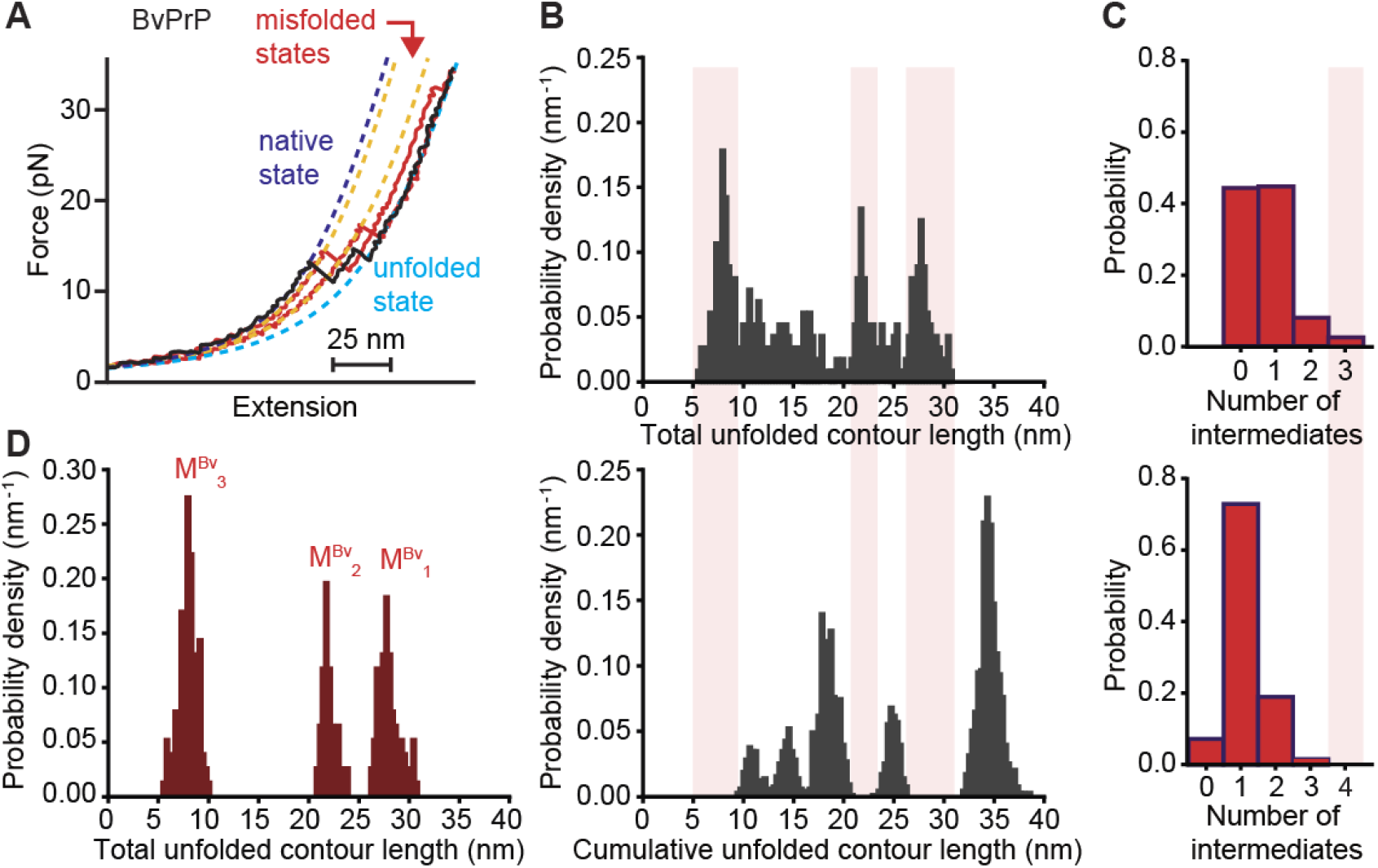
Misfolding of BvPrP. (A) Over ¼ of BvPrP unfolding FECs show non-native total Δ*L*_c_ (red) instead of native Δ*L*_c_ (black). Dashed lines: WLC fits. (B) Distribution of unfolded *L*_c_ in initial state for unfolding FECs with non-native total Δ*L*_c_ (top) compared to unfolded *L*_c_ distribution for all states in native unfolding (bottom). Pink bars: lengths not matching anything in the native distribution, implying misfolded states. (C) Number of intermediates in FECs unfolding from non-native (top) and native (bottom) initial states. Pink bar: FECs with non-native number of intermediates, implying misfolding. (D) Distribution of total Δ*L*_c_ for FECs identified as misfolded, indicating at least three distinct misfolded states.

To test if such FECs reflected misfolded states or intermediates on the native pathway, we examined the distribution of total unfolded contour lengths for the initial states in all unfolding FECs showing a total Δ*L*_c_ shorter than native. We compared this distribution (Fig. 5B, top) with the distribution of cumulative unfolded contour lengths for intermediates in native unfolding (Fig. 5B, bottom). We found that some of the FECs with shorter-than-native total Δ*L*_c_ matched lengths found in the native-intermediate distribution, indicating that in these cases the protein might have been in a partially folded native intermediate. However, a significant number of the FECs had total unfolded *L*_c_ that did not align with any of the intermediate states on the native pathway. Indeed, we distinguished three such peaks from this distribution: at 8 nm, 22 nm, and 28 nm (Fig. 5B, pink). These states were not consistent with any intermediate on the native pathway and hence presumably reflected misfolding of the protein.

An additional criterion for detecting misfolding is based on the number of intermediates seen in the FECs: given that native unfolding never showed more than 3 intermediates in total (Fig. 5C, bottom), any unfolding FEC with shorter-than-native total Δ*L*_c_ that starts from an intermediate on the native pathway should thus show at most two intermediates. However, a few of the FECs did, in fact, show three intermediates (Fig. 5C, top), implying that they started from a state not on the native pathway—a misfolded state. The misfolded states identified in this way matched the same total Δ*L*_c_ values found using the length analysis above. We denoted the three misfolded states found through these analyses as M^Bv^_1_ at total Δ*L*_c_ = 28 nm, M^Bv^_2_ at 22 nm, and M^Bv^_3_ at 8 nm (Fig. 5D, Table S3). Overall, ∼19% of all BvPrP unfolding FECs showed evidence that BvPrP started off in a misfolded state.

Finally, to explore the pathways for misfolding and how they differed from the native folding pathways, we examined the refolding FECs that were measured just prior to the unfolding FECs that started from a misfolded state—*i*.*e*., the subset of refolding curves in which the protein was undergoing misfolding. Plotting the distribution of cumulative unfolded contour lengths based on the transitions from the unfolded state to the misfolded states (Fig. 6A), by analogy to the situation above for native folding, the peaks in this distribution reflect intermediates on the misfolding pathway. For refolding FECs ending up in M^Bv^_1_ (Fig. 6A, top), there was no evidence of any intermediates: the protein always jumped directly from the unfolded state to M^Bv^_1_. For refolding FECs ending at M^Bv^_2_, one intermediate state was observed, at the same length as M^Bv^_1_. For curves ending at M^Bv^_3_, three intermediate states were observed, of which two had the same lengths as M^Bv^_1_ and M^Bv^_2_. The similarity of the lengths for many of these states could reflect the presence of a single pathway connecting multiple distinct misfolded states that are sufficiently long-lived to persist for many seconds during FEC measurements (Fig. 6B), or multiple pathways featuring misfolded states that just coincidentally share the same lengths.

**Fig. 6:**
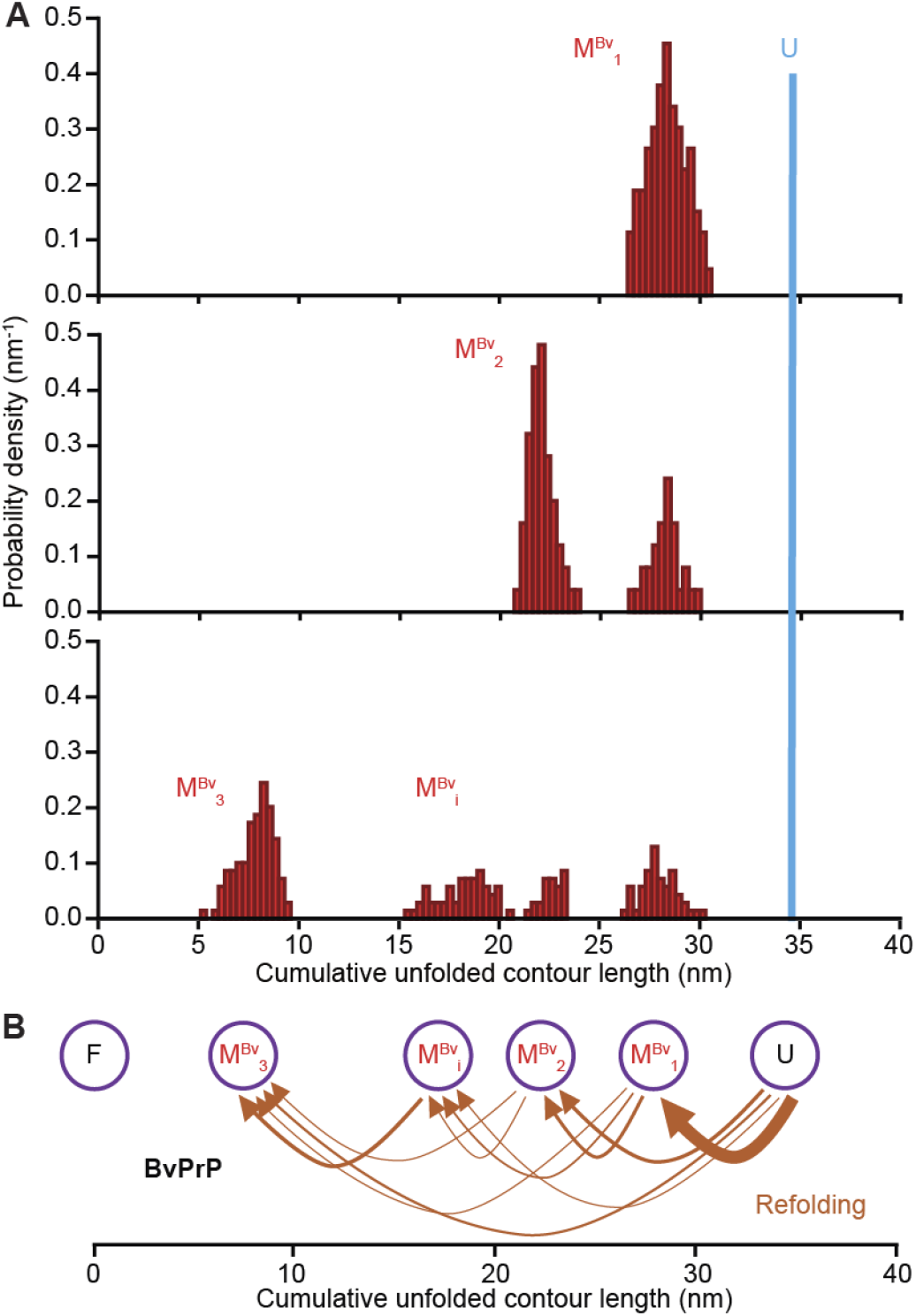
BvPrP misfolding pathway. (A) Distribution of unfolded *L*_c_ for all states in refolding FECs leading to misfolded states M^Bv^_1_ (top), M^Bv^_2_ (middle), and M^Bv^_3_ (bottom). M^Bv^_1_ may act as an intermediate leading to M^Bv^_2_, and M^Bv^_2_ along with M^Bv^_i_ may do the same leading to M^Bv^_3_. Blue line indicates position of unfolded state. (B) Transition map for BvPrP misfolding, assuming a single pathway. Arrow thickness proportional to relative frequency of transition.

## DISCUSSION

Here we have examined the folding of PrP from three different species (two rodents and one canid) under the same conditions at the single-molecule level. We observed profoundly different behavior between the three species, despite the fact that they all form the same fold: different levels of cooperativity in folding, different intermediates on the folding pathways, and qualitatively different energy barriers (homogeneous vs heterogeneous). These differences are particularly remarkable given the very high level of sequence identity and even higher level of sequence similarity: there are only 2 non-conservative amino-acid differences between HaPrP and BvPrP in the 104-amino-acid structured domain, and only 7 or 8 between CaPrP and the rodent PrPs. Furthermore, the two rodent PrPs are evolutionarily very close,^12^ yet nevertheless display dramatic differences in their folding dynamics, implying quite different folding mechanisms. In stark contrast, previous work has shown that key features of folding mechanisms such as transition states and intermediates can be conserved across homologs with considerably lower sequence similarity and over very long evolutionary timescales.^64–66^

Given that only a few mutations are sufficient to cause significant changes in the barriers and intermediates involved in PrP folding, the energy landscape for PrP must be very sensitive to small perturbations, such that subtle changes in the intramolecular energies redirect the folding to different pathways with different barriers. This picture is consistent with previous work finding that the native structure of PrP is highly frustrated,^67^ with many unsatisfied interactions that could in principle be turned to stabilize alternative structures. Although initially proposed to explain the misfolding propensity of PrP, such frustration could also explain the low robustness of the native folding pathway to mutation seen here, further supporting the notion that a peculiar fragility of the energy landscape for PrP folding is a key feature underlying this protein’s propensity to misfold and propagate its misfolding infectiously. Intriguingly, PrP is proposed to have evolved from a membrane protein with poor solubility, which could account for its frustrated landscape.^68,69^

Looking at the sequence differences between CaPrP, HaPrP, and BvPrP to see what might explain the different folding behavior, we note that for CaPrP, D159 (replacing N in the rodent PrPs) has been identified as a crucial residue conferring resistance in dogs, possibly by generating a salt bridge with R135 in the loop linking strand 1 and helix 1 or electrostatic interactions with nearby positively charged residues.^17^ These interactions could also stabilize the proposed structure for intermediate I^Ca^_3_ (folded helix 1 and strands 1/2), which is also a component of the structure in I^Ca^_1_ and I^Ca^_2_ but not seen in any intermediates for the rodent PrPs. Given that helix 1 has been proposed to be a locus of misfolding,^70,71^ the early native-like folding of helix 1 in CaPrP might confer protection against misfolding and thus help explain canid resistance to prion diseases. Turning to a comparison of the rodent PrPs, the significant differences in their folding (*e*.*g*. two-state versus multi-state) are harder to explain, as there are only two non-conservative mutations, at residues 215 in helix 3 (T in HaPrP, V in BvPrP) and 230 at the C terminus (R in HaPrP, S in BvPrP), alongside 4 conservative mutations (M139I, I203V, I205M, D227E). We speculate that the resulting changes in electrostatic interactions between the C terminus and the rigid loop, and/or solvent interactions in helix 3, may account for the differences in folding.

The most dramatic difference in the folding of the three PrPs studied here is the detection of long-lived misfolded states in BvPrP. Notably, these misfolded states are monomeric. Misfolded states were observed in single HaPrP monomers previously, but they were very short-lived,^48^ in line with the assumption that misfolded PrP is generally oligomeric.^72^ Although long-lived misfolded monomers have been reported under certain conditions in ensemble assays,^73^ stable misfolding at the single-molecule level has only ever been seen previously in oligomers or aggregates.^50–53^ The observation of long-lived misfolded states in BvPrP monomers presents an opportunity for studying misfolding mechanisms of PrP in more detail. Intriguingly, the misfolded states observed in BvPrP do not appear to involve any partially-native structure: they form directly from the unfolded state without branching off the native pathway, unlike the misfolding mechanisms observed in several other proteins.^40,41^ In particular, we see no evidence for misfolding arising from a partially native intermediate as proposed previously for human PrP.^74^ Such a result makes sense in the light of the structural models proposed for PrP^Sc^, in which the native helical content is completely removed:^75,76^ all native intermediates contain helices that would need to be unfolded to ensure a purely β-structured conformation. The misfolding mechanism thus appears to be entirely independent of native folding. This result reinforces the view that unfolded states play a crucial role in PrP misfolding.^37,54,55,77,78^ Intriguingly, the lengths for M^Bv^_2_, and M^Bv^_i_ match those for two of the frequently occurring short-lived misfolded states detected previously in HaPrP,^48^ suggesting that they might be similar structurally.

Finally, turning to the question of what features of the folding might relate to differences in misfolding and disease susceptibility, we note that folding cooperativity is clearly uncorrelated with either—the presence of more intermediate states does not induce increased susceptibility, as HaPrP (intermediate susceptibility) is the most cooperative whereas CaPrP (resistant) is the least. Differences in the native folding/unfolding pathways also seem to be unrelated to susceptibility or misfolding propensity. Although no quantitative trends in the unloaded unfolding rates or barrier heights correlating with susceptibility could be discerned, qualitatively the amount of hysteresis in folding/unfolding increased with susceptibility, suggesting that slower folding rates and/or higher barriers (which produce hysteresis) could help enhance susceptibility. We speculate that higher barriers/slower refolding may increase the tendency to misfold by providing more time for the protein to find its way onto misfolding pathways. Presumably, the high susceptibility of BvPrP is related to the observation that it is also the most likely to misfold, and can sustain longlived misfolded states in monomers.

## METHODS

### Sample preparation

Samples of PrP from each species, with sequence truncated to residues 90-231 (forming the protease-resistant core of PrP^Sc^), were expressed recombinantly in *E. coli*, purified, and refolded as described previously.^48^ In each construct, a cysteine residue was added for handle attachments at both termini. Handles were attached by first reducing the protein with tris(2-carboxyethyl)phosphine (TCEP), removing excess TCEP by spin filtration (Amicon Ultracel-10K 0.5 ml centrifugal filters), and incubating the protein in 100-fold molar excess of 2,2′-dithiodipyridine (DTDP) at 4 °C for 24 h to activate the cysteines. Excess DTDP was then removed by spin filtration, and the protein was incubated at 4 °C for 24 h with sulfhydryl-labelled DNA handles (one of length 802 bp labelled with biotin, the other of length 2100 bp labelled with digoxigenin). We used pyridine-2-thione release and mass spectrometry to assess that the PrP was labelled with either one or two DTDP molecules (average of 1.5–1.6). The resulting protein-DNA constructs were incubated at ∼100 pM with 250 pM polystyrene beads (600-nm diameter functionalized with avidin, 820-nm diameter functionalized with anti-digoxigenin) to form dumbbells. Dumbbells were diluted to ∼500 fM in measurement buffer (50 mM MOPS, pH 7.0, 200 mM KCl, along with an oxygen scavenging system containing 8mU/μl glucose oxidase, 20mU/μl catalase, and 0.01% w/v D-glucose) and placed in a sample cell for optical trapping.

### SMFS measurements

SMFS measurements were made using custom-built optical tweezers following the procedure described previously.^45^ The traps were moved apart in 1–2 nm increments at a pulling speed of 125–250 nm/s to produce FECs. Data was sampled at 20kHz, filtered online with an eight-pole Bessel filter (Krohn-Hite) at the Nyquist frequency. The stiffness of the traps was calibrated to 0.37 and 0.56 pN/nm. We waited for 2–15s at near-zero force between successive pulls to allow the protein to refold.

### Contour length analysis

The change in the contour length (Δ*L*_c_) during unfolding and refolding transitions was quantified by fitting the FECs to an extensible WLC model as done previously:^45,48^

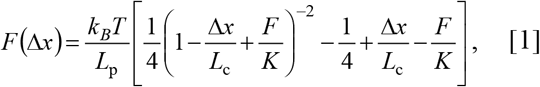

where *L*_p_ is the polymer persistence length, *L*_c_ the contour length, and *K* the elastic modulus. Two WLCs were used in series, one for the handles and the other for the unfolded protein. We treated *L*_c_, *L*_p_ and *K* for the DNA handles as free parameters when fitting the completely folded segment of the FECs, then treated these parameters as fixed when fitting intermediate or unfolded segments of the FECs. *L*_p_ and *K* for the protein were also treated as fixed in these fits, with *L*_p_ = 0.85 nm and *K* = 2000 pN,^45^ so that *L*_c_ was the only free parameter for fitting intermediate and unfolded states.

The contour length changes (Δ*L*_c_) obtained from fitting each branch of the FECs were used to determine the distribution of cumulative Δ*L*_c_ starting from the folded state as described previously.^41^ The cumulative Δ*L*_c_ distributions were similar for each molecule measured for a given PrP variant, indicating all molecules behaved similarly. The results from all molecules were therefore combined into a single histogram for analysis. The contour length change (Δ*L*_c_) predicted for complete unfolding from the NMR structures of the PrP variants was found as Δ*L*_c_ = *n*_aa_×*L*c^aa^ − *d*_T_, where *n*_aa_ is the number of structured amino acids, *L*_c_^aa^ =0.36 nm/aa is the crystallographic contour length of an amino acid, and *d*_T_ is the distance between the termini.

### Unfolding force distribution analysis

Unfolding force distributions, *p*(*F*), were fit to models for escape over a barrier that is either homogeneous^60^ or heterogeneous.^61^ Assuming a linear-cubic potential, the first model yields:

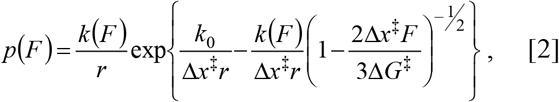

Where

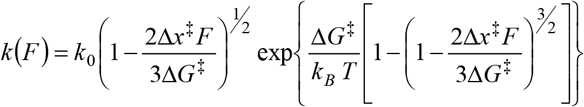

is the force-depending unfolding rate, *k*_0_ is the unfolding rate at zero force, Δ*x*^‡^ is the distance from the folded state to the energy barrier for unfolding (transition state), Δ*G*^‡^ is the barrier height, and *r* is the loading rate. The second model assumes conversion between heterogeneous energy barriers, and gives

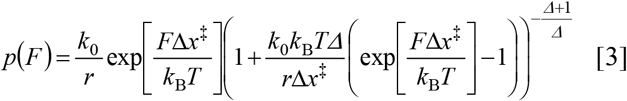

where *k*_0_ is the zero-force unfolding rate averaged over the ensemble of initial states, Δ*x*^‡^ is the average distance to the transition state from the ensemble of starting states, and *Δ* is a dimensionless parameter reflecting the extent of heterogeneity (where larger *Δ* indicates greater heterogeneity).

### Network analysis of possible intermediate structures and folding pathways

We used the approach described previously^41^ to determine possible structures of intermediate states. We first identified all possible substructures of the native fold of PrP consisting of some subset of structural elements from the native fold, assuming that (a) loops did not provide any mechanical resistance and hence were excluded unless sequestered between folded strands and/or helices, and (b) the folded elements within each substructure retained native-like structure. For each substructure, we then calculated the expected cumulative contour length of unfolded protein, *L*_c_, via:

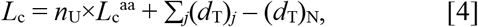

where *n*_U_ is the number of unfolded residues in the substructure, (*d*_T_)_N_ is the distance between the termini of the native structure, and Σ_j_(*d*_T_)_j_ is the sum over the distances between the termini of each discrete structured region in the intermediate (where more than one structured region exists; for example, owing to an unfolded loop between two folded helices). The possible intermediates are listed with their expected *L*_c_ values in Table S2. For each intermediate in a FEC, we compared the measured *L*_c_ values to those expected for all possible substructures to identify prospective matches, assuming a match if they were the same within 2 nm (to account for uncertainty). This approach allows for multiple distinct substructures as matches. The matching substructures were then employed as nodes in a network analysis to identify all feasible structural pathways through the sequence of intermediates observed in each given FEC (Fig. S4), under the assumption that structural elements did not refold once unfolded (and vice versa). Finally, we applied this analysis to all FECs for a given PrP variant to find the minimal set of intermediate structures and pathways through them that explained the maximum number of FECs.

### MD simulations of PrP substructures

To check if the substructures proposed as intermediates are plausibly stable structures when isolated from within the native fold, we performed all-atom MD simulations using Amber 18.^79^ Substructures were prepared for simulations by truncating the PDB structures for each PrP variant (CaPrP: 1XYK, HaPrP: 1B10, BvPrP: 2K56) based on the residues listed in Table S2, and capping the termini of each substructure with a neutral acetyl capping group (−C(=O)−CH3) and a neutral methylamine capping group (C(=O)−NH−CH3) at the N- and C-termini, respectively. We used the ff14SB force field to parameterize the protein and each sub-structure was solvated in TIP3P explicit water. Na^+^ and Cl^−^ ions were added at a concentration of 0.15 M. For each substructure, three 0.75-µs long simulations were performed at 310 K with heavy restraints on the cap residues. Using the CPPTRAJ module in Amber, the structures from the last 250 ns from all three replicate trajectories for each substructure were pooled together to compute the secondary structure assignment, based on the dominant secondary structure type for each residue. Substructures were considered unstable and rejected as possibilities for the pathway analysis if over 25% of their secondary structure content unfolded during the simulations.

## Conflict of interest

The authors declare no conflict.

## Author contributions

UA and SP measured data; UA, SP, and RVS analysed data; CRG, UA, and SP prepared reagents; RVS performed simulations; MTW, UA, SP, and RVS wrote the paper; all authors edited the paper.

## Acknowledgements

This work was supported by the Alberta Prion Research Institute (grant reference numbers 201600014 and 201800008) and the Natural Sciences and Engineering Research Council of Canada (grant reference number RGPIN-2018-04673). UA acknowledges fellowship support from Alberta Innovates.

## Supplementary information

**Table S1:** Contour length changes for native folding/unfolding. The Δ*L*_c_ observed for unfolded and intermediate states with respect to the native state for CaPrP (total of 1,320 FECs from 7 molecules), HaPrP (total of 3,250 FECs from 7 molecules), and BvPrP (total of 2,700 FECs from 11 molecules). Uncertainty represents standard error on the mean value (averaged across all molecules).

| Species | Observed unfolded state (nm) | Intermediates | Observed unfolding $\Delta L_c$ (nm) | Observed refolding $\Delta L_c$ (nm) |
| --- | --- | --- | --- | --- |
| Dog | $34.9 \pm 0.3$ | $I_{Ca_1}$ | $12.2 \pm 0.6$ | $13.0 \pm 0.5$ |
| | | $I_{Ca_2}$ | $18.2 \pm 0.4$ | $18.1 \pm 0.4$ |
| | | $I_{Ca_3}$ | $22.5 \pm 0.3$ | $22.7 \pm 0.4$ |
| | | $I_{Ca_4}$ | $27.8 \pm 0.4$ | $27.4 \pm 0.3$ |
| Hamster | $34.1 \pm 0.4$ | N/A | N/A | N/A |
| Bank vole | $34.5 \pm 0.3$ | $I_{Bv_1}$ | $10.7 \pm 0.2$ | $10.6 \pm 0.3$ |
| | | $I_{Bv_2}$ | $14.1 \pm 0.2$ | $14.2 \pm 0.2$ |
| | | $I_{Bv_3}$ | $18.2 \pm 0.3$ | $18.5 \pm 0.3$ |
| | | $I_{Bv_4}$ | $24.9 \pm 0.2$ | $25.2 \pm 0.2$ |

**Table S2:**
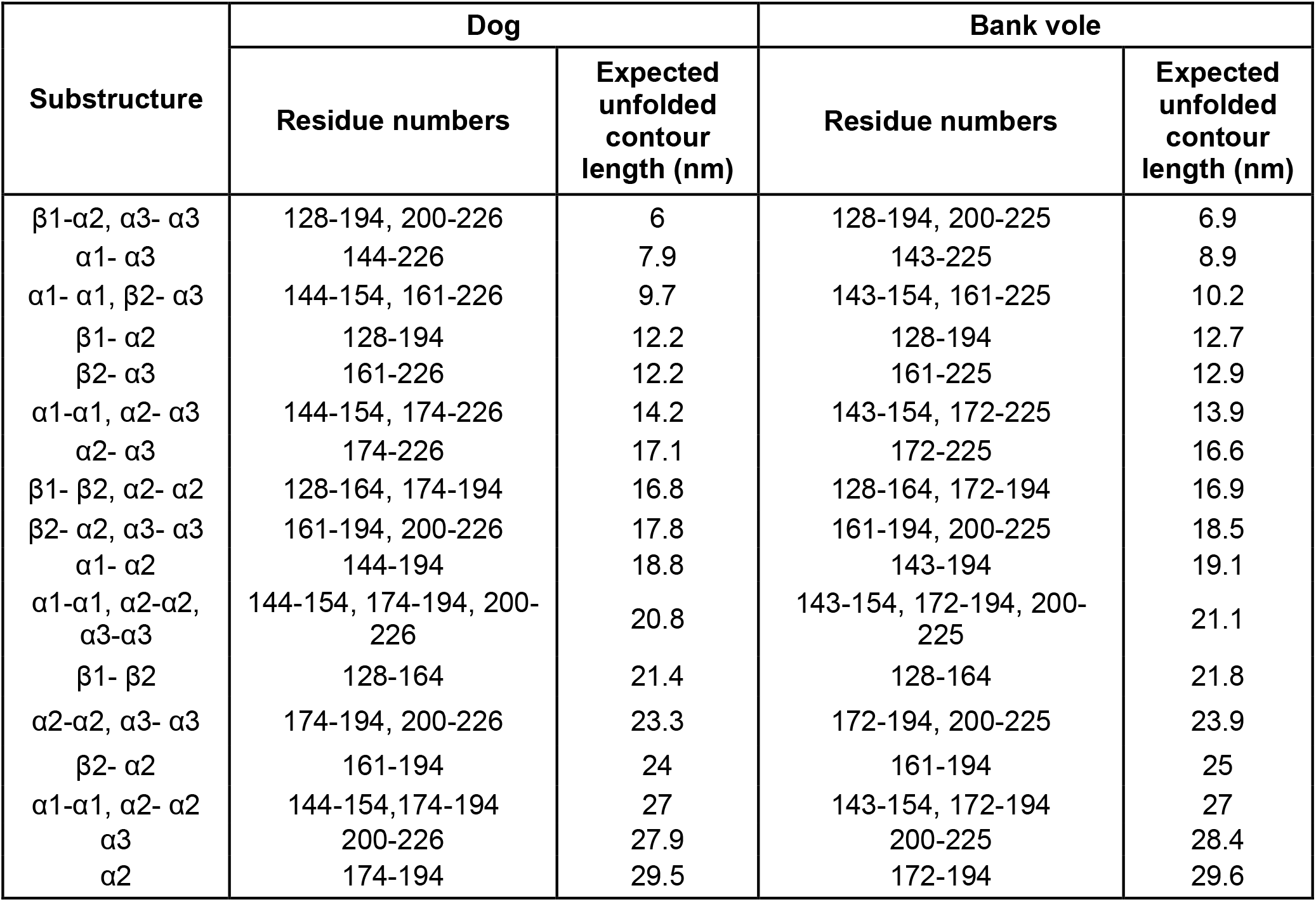
Possible structures of intermediate states. Sub-structures of the native fold (consisting of different combinations of secondary structural elements within the native fold) hypothesized as possible intermediate states, along with the corresponding expected contour length of unfolded protein.

| Substructure | Dog |  | Bank vole |  |
| --- | --- | --- | --- | --- |
|  | Residue numbers | Expected unfolded contour length (nm) | Residue numbers | Expected unfolded contour length (nm) |
| $\beta 1$-$\alpha 2$, $\alpha 3$-$\alpha 3$ | 128-194, 200-226 | 6 | 128-194, 200-225 | 6.9 |
| $\alpha 1$-$\alpha 3$ | 144-226 | 7.9 | 143-225 | 8.9 |
| $\alpha 1$-$\alpha 1$, $\beta 2$-$\alpha 3$ | 144-154, 161-226 | 9.7 | 143-154, 161-225 | 10.2 |
| $\beta 1$-$\alpha 2$ | 128-194 | 12.2 | 128-194 | 12.7 |
| $\beta 2$-$\alpha 3$ | 161-226 | 12.2 | 161-225 | 12.9 |
| $\alpha 1$-$\alpha 1$, $\alpha 2$-$\alpha 3$ | 144-154, 174-226 | 14.2 | 143-154, 172-225 | 13.9 |
| $\alpha 2$-$\alpha 3$ | 174-226 | 17.1 | 172-225 | 16.6 |
| $\beta 1$-$\beta 2$, $\alpha 2$-$\alpha 2$ | 128-164, 174-194 | 16.8 | 128-164, 172-194 | 16.9 |
| $\beta 2$-$\alpha 2$, $\alpha 3$-$\alpha 3$ | 161-194, 200-226 | 17.8 | 161-194, 200-225 | 18.5 |
| $\alpha 1$-$\alpha 2$ | 144-194 | 18.8 | 143-194 | 19.1 |
| $\alpha 1$-$\alpha 1$, $\alpha 2$-$\alpha 2$, $\alpha 3$-$\alpha 3$ | 144-154, 174-194, 200-226 | 20.8 | 143-154, 172-194, 200-225 | 21.1 |
| $\beta 1$-$\beta 2$ | 128-164 | 21.4 | 128-164 | 21.8 |
| $\alpha 2$-$\alpha 2$, $\alpha 3$-$\alpha 3$ | 174-194, 200-226 | 23.3 | 172-194, 200-225 | 23.9 |
| $\beta 2$-$\alpha 2$ | 161-194 | 24 | 161-194 | 25 |
| $\alpha 1$-$\alpha 1$, $\alpha 2$-$\alpha 2$ | 144-154, 174-194 | 27 | 143-154, 172-194 | 27 |
| $\alpha 3$ | 200-226 | 27.9 | 200-225 | 28.4 |
| $\alpha 2$ | 174-194 | 29.5 | 172-194 | 29.6 |

**Table S3:** Misfolded states in BvPrP. Observed contour lengths of unfolded protein in misfolded states. Uncertainty represents standard error on the mean value (averaged across all molecules).

| Misfolded states | Total unfolded contour length (nm) |
| --- | --- |
| $M^{Bv_1}$ | $28.0 \pm 0.3$ |
| $M^{Bv_2}$ | $22.0 \pm 0.2$ |
| $M^{Bv_3}$ | $8.0 \pm 0.3$ |

**Fig. S1:**
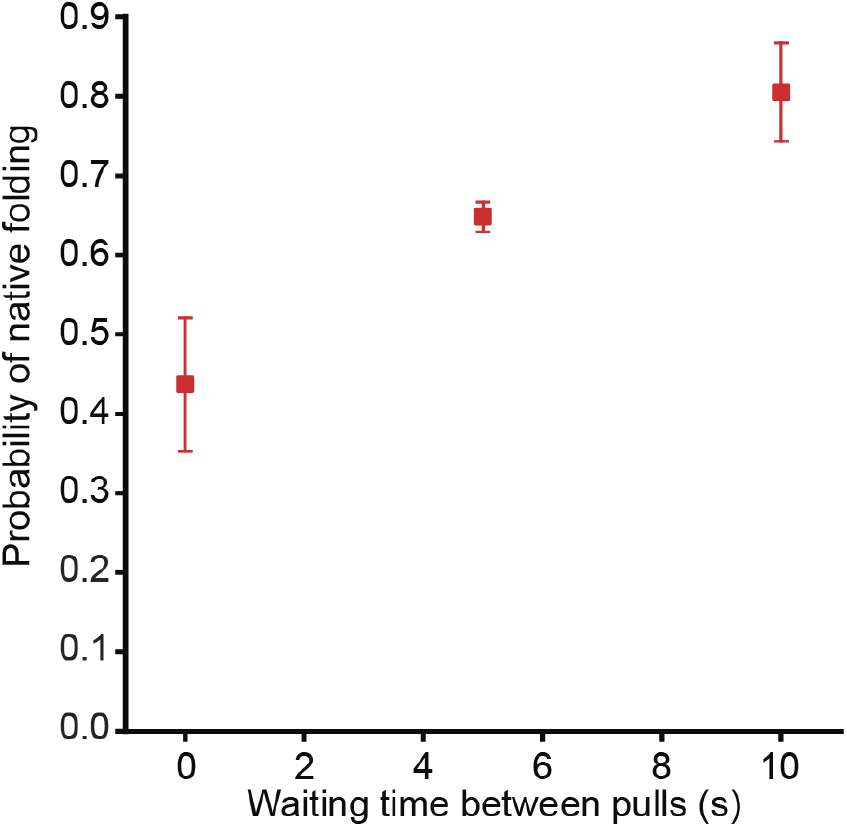
Slow refolding of BvPrP. The number of native full-length FECs increased as the waiting time between pulls (near zero force) increased.

**Fig. S2:**
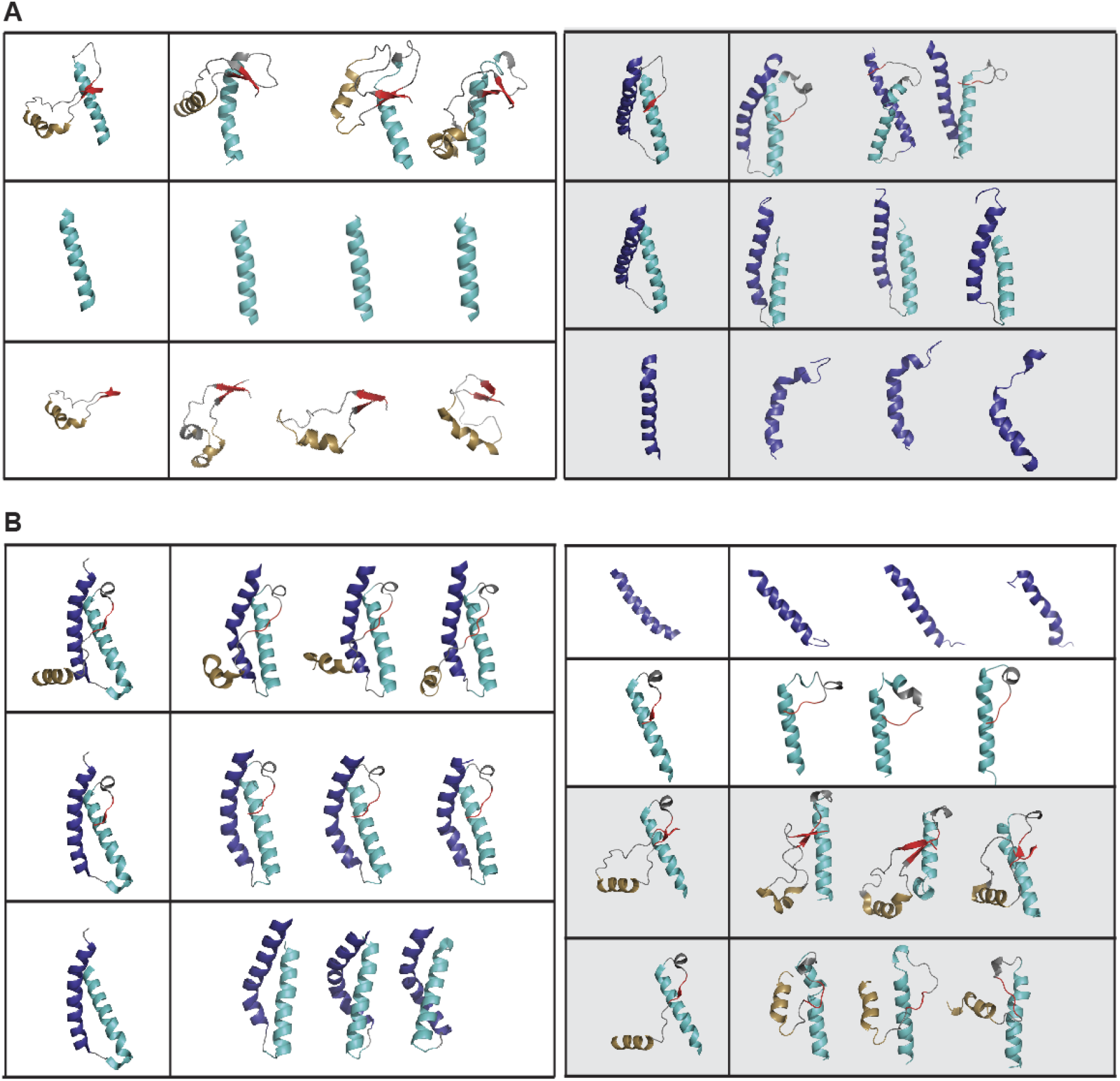
MD simulations of possible substructures. Possible substructures for (A) CaPrP and (B) BvPrP were tested for stability in MD trajectories. In each case, the substructure at the start of the simulation is shown on the left, and the final structures from three MD simulations are shown on the right. Secondary structure representations for each residue are assigned based on the dominant secondary structure of the residues in the last 250 ns of the trajectories (α-helix, 3-10 helix, π-helix, and turns are represented as helices; parallel and anti-parallel β-sheets are represented as sheets; bends and no secondary structure are represented as loops). Grey panels represent substructures for alternate pathways shown in Fig. S3.

**Fig. S3:**
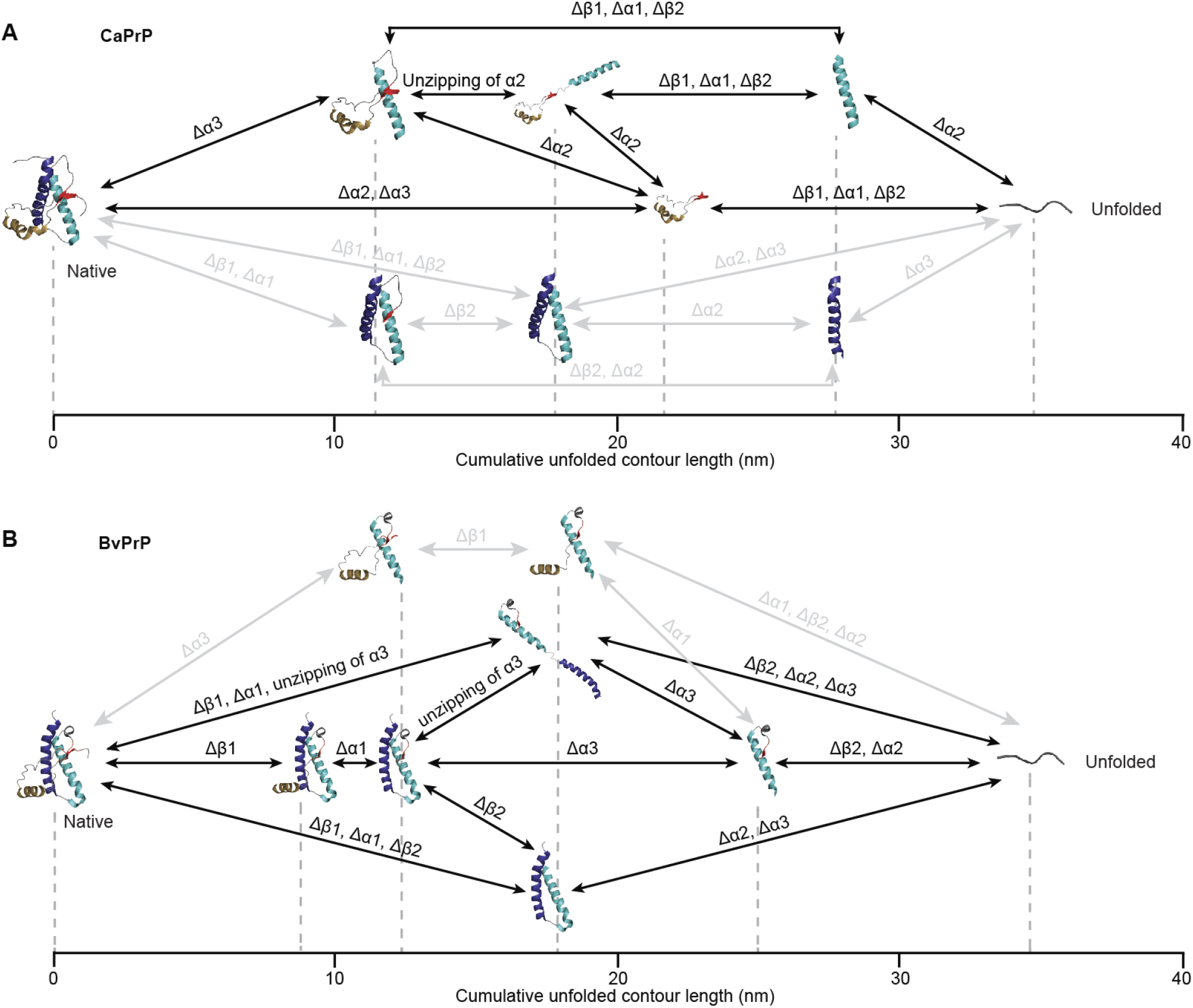
Alternative native folding pathways. Alternative pathways for native folding of (A) CaPrP and (B) BvPrP are shown (grey) along with the dominant pathways. Adding these extra pathways increased the explanatory power of the model marginally, accounting for ∼90% of FECs in CaPrP and ∼89% of FECs in BvPrP.

**Fig. S4:**
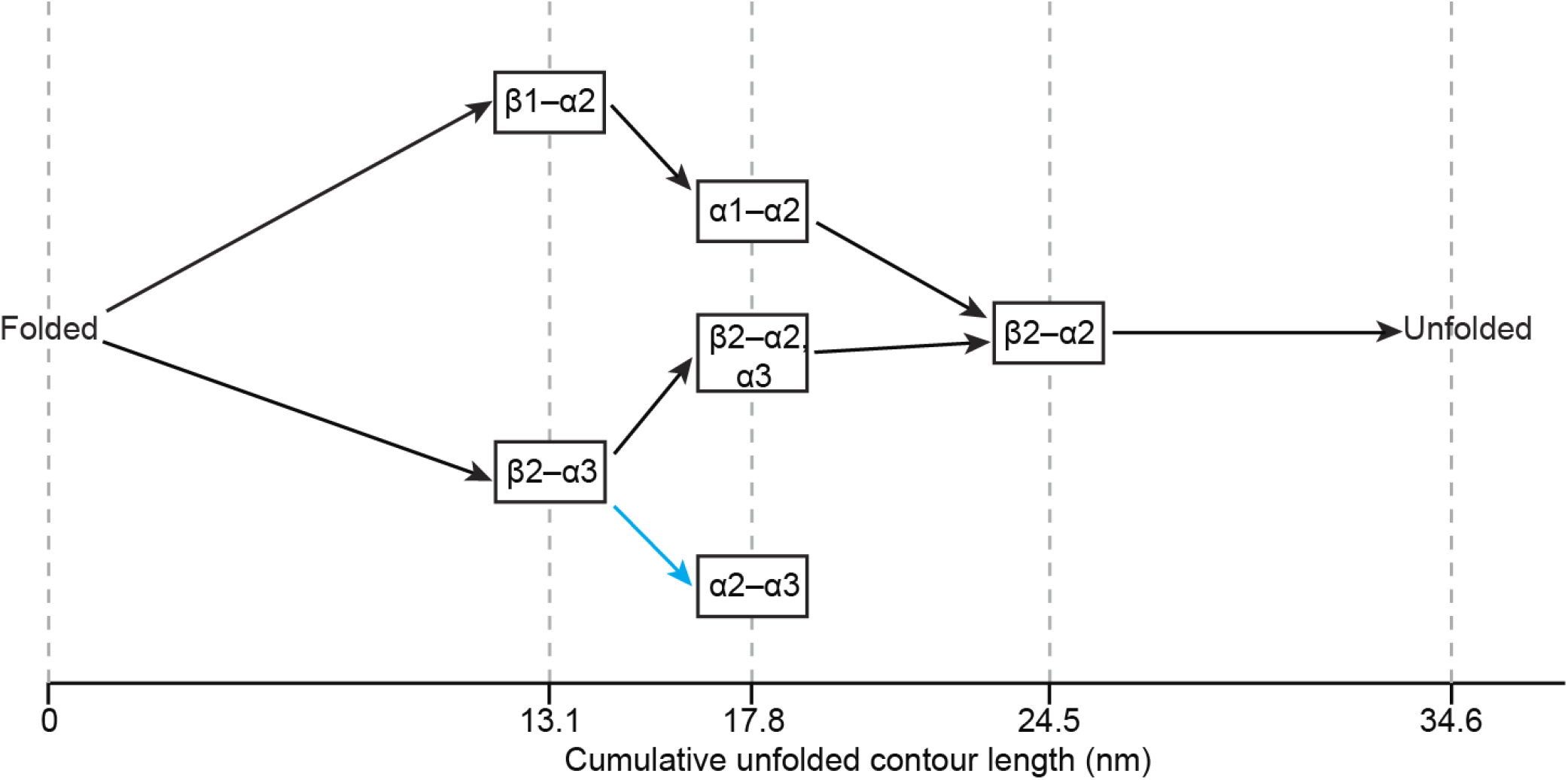
Representative network analysis to identify potential sub-structures along the folding pathway from a FEC. Displaying the experimentally observed cumulative Δ*L*_c_ values for a particular FEC as dashed lines, all the possible sub-structures consistent with these values are depicted, labeled with the portion of the structure that remains folded. Two viable pathways from folded to unfolded states for this FEC are shown as black arrows; one non-productive pathway (cyan) is unable to connect to the unfolded structure, because α2-α3 cannot lead to β2-α2 without simultaneous refolding of β2 and unfolding of α3.

## Notes

### Competing Interest Statement

The authors have declared no competing interest.

## References

1. Cobb, N. J. & Surewicz, W. K. Prion diseases and their biochemical mechanisms. Biochemistry 48, 2574–2585 (2009).

2. Colby, D. W. & Prusiner, S. B. Prions. Cold Spring Harb. Perspect. Biol. 3, 1–22 (2011).

3. Baldwin, M. A. et al. Spectroscopic characterization of conformational differences between PrPC and PrPSc: an α-helix to β-sheet transition. Philos. Trans. R. Soc. Lond. B Biol. Sci. 343, 435–441 (1994).

4. Caughey, B. W. et al. Secondary structure analysis of the scrapie-associated protein PrP 27–30 in water by infrared spectroscopy. Biochemistry 30, 7672–7680 (1991).

5. Gasset, M. et al. Predicted alpha-helical regions of the prion protein when synthesized as peptides form amyloid. Proc. Natl. Acad. Sci. USA 89, 10940–10944 (1992).

6. Pan, K. M. et al. Conversion of α-helices into β-sheets features in the formation of the scrapie prion proteins. Proc. Natl. Acad. Sci. USA 90, 10962–10966 (1993).

7. Caughey, B., Baron, G. S., Chesebro, B. & Jeffrey, M. Getting a grip on prions: Oligomers, amyloids, and pathological membrane interactions. Annu. Rev. Biochem. 78, 177–204 (2009).

8. McAlary, L., Plotkin, S. S., Yerbury, J. J. & Cashman, N. R. Prion-like propagation of protein misfolding and aggregation in amyotrophic lateral sclerosis. Front. Mol. Neurosci. 12, 262 (2019).

9. Vaquer-Alicea, J. & Diamond, M. I. Propagation of protein aggregation in neurodegenerative diseases. Annu. Rev. Biochem. 88, 785–810 (2019).

10. Soto, C. & Pritzkow, S. Protein misfolding, aggregation, and conformational strains in neurodegenerative diseases. Nat. Neurosci. 21, 1332–1340 (2018).

11. Jucker, M. & Walker, L. C. Propagation and spread of pathogenic protein assemblies in neurodegenerative diseases. Nat. Neurosci. 21, 1341–1349 (2018).

12. Cortez, L. M. et al. Probing the origin of prion protein misfolding via reconstruction of ancestral proteins. Protein Science 31, e4477 (2022).

13. Wopfner, F. et al. Analysis of 27 mammalian and 9 avian PrPs reveals high conservation of flexible regions of the prion protein. J. Mol. Biol. 289, 1163–1178 (1999).

14. Schätzl, H. M., Costa, M. Da, Taylor, L., Cohen, F. E. & Prusiner, S. B. Prion protein gene variation among primates. J. Mol. Biol. 245, 362–374 (1995).

15. Kirkwood, J. K. & Cunningham, A. A. Epidemiological observations on spongiform encephalopathies in captive wild animals in the British Isles. Vet. Rec. 135, 296–303 (1994).

16. Bian, J. et al. Prion replication without host adaptation during interspecies transmissions. Proc. Natl. Acad. Sci. USA 114, 1141–1146 (2017).

17. Fernández-Borges, N. et al. Unraveling the key to the resistance of canids to prion diseases. PLoS Pathog. 13, e1006716 (2017).

18. Barlow, R. M. & Rennie, J. C. The fate of ME7 scrapie infection in rats, guinea-pigs and rabbits. Res. Vet. Sci. 21, 110–111 (1976).

19. Hammarström, P. & Nyström, S. Porcine prion protein amyloid. Prion 9, 266–277 (2015).

20. Vorberg, I., Groschup, M. H., Pfaff, E. & Priola, S. A. Multiple amino acid residues within the rabbit prion protein inhibit formation of its abnormal isoform. J. Virol. 77, 2003–2009 (2003).

21. Fernandez-Funez, P. et al. Pulling rabbits to reveal the secrets of the prion protein. Commun. Integr. Biol. 4, 262–266 (2011).

22. Gibbs, C. J. & Carleton Gajdusek, D. Experimental subacute spongiform virus encephalopathies in primates and other laboratory animals. Science 182, 67–68 (1973).

23. Ishida, Y. et al. Association of chronic wasting disease susceptibility with prion protein variation in white-tailed deer (Odocoileus virginianus). Prion 14, 214–225 (2020).

24. Nonno, R. et al. Efficient transmission and characterization of Creutzfeldt–Jakob disease strains in bank voles. PLoS Pathog. 2, e12 (2006).

25. Prusiner, S. B. et al. Transgenetic studies implicate interactions between homologous PrP isoforms in scrapie prion replication. Cell 63, 673–686 (1990).

26. Sarradin, P. et al. Transgenic rabbits expressing ovine PrP are susceptible to scrapie. PLoS Pathog. 11, e1005077 (2015).

27. Wen, Y. et al. Unique structural characteristics of the rabbit prion protein. J. Biol. Chem. 285, 31682–31693 (2010).

28. Lysek, D. A. et al. Prion protein NMR structures of cats, dogs, pigs, and sheep. Proc. Natl. Acad. Sci. USA 102, 640–645 (2005).

29. James, T. L. et al. Solution structure of a 142-residue recombinant prion protein corresponding to the infectious fragment of the scrapie isoform. Proc. Natl. Acad. Sci. USA 94, 10086–10091 (1997).

30. Christen, B., Pérez, D. R., Hornemann, S. & Wüthrich, K. NMR structure of the bank vole prion protein at 20 °C contains a structured loop of residues 165–171. J. Mol. Biol. 383, 306–312 (2008).

31. Agrimi, U., Nonno, R., Dell’Omo, G., Bari, D. & Conte, M. A. Prion protein amino acid determinants of differential susceptibility and molecular feature of prion strains in mice and voles. PLoS Pathog. 4, e1000113 (2008).

32. Asante, E. A. et al. A naturally occurring variant of the human prion protein completely prevents prion disease. Nature 522, 478–481 (2015).

33. Baral, P. K., Swayampakula, M., Aguzzi, A. & James, M. G. X-ray structural and molecular dynamical studies of the globular domains of cow, deer, elk and Syrian hamster prion proteins. J. Struct. Biol. 192, 37–47 (2015).

34. Christen, B., Damberger, F. F., Pérez, D. R., Hornemann, S. & Wüthrich, K. Structural plasticity of the cellular prion protein and implications in health and disease. Proc. Natl. Acad. Sci. USA 110, 8549–8554 (2013).

35. Hasegawa, K., Mohri, S. & Yokoyama, T. Comparison of the local structural stabilities of mammalian prion protein (PrP) by fragment molecular orbital calculations. Prion 7, 185–191 (2013).

36. Qasim Khan, M. et al. Prion disease susceptibility is affected by β-structure folding propensity and local side-chain interactions in PrP. Proc. Natl. Acad. Sci. USA 107, 19808–19813 (2010).

37. Srivastava, K. R. & Lapidus, L. J. Prion protein dynamics before aggregation. Proc. Natl. Acad. Sci. USA 114, 3572–3577 (2017).

38. Greenleaf, W. J., Woodside, M. T. & Block, S. M. High-resolution, single-molecule measurements of biomolecular motion. Annu. Rev. Biophys. Biomol. Struct. 36, 171–190 (2007).

39. Hoffmann, A., Neupane, K. & Woodside, M. T. Single-molecule assays for investigating protein misfolding and aggregation. Physical Chemistry Chemical Physics 15, 7934–7948 (2013).

40. Stigler, J., Ziegler, F., Gieseke, A., Gebhardt, J. C. M. & Rief, M. The complex folding network of single cal-modulin molecules. Science 334, 512–516 (2011).

41. Sen Mojumdar, S. et al. Partially native intermediates mediate misfolding of SOD1 in single-molecule folding trajectories. Nat. Commun. 8, 1881 (2017).

42. Ferreon, A. C. M., Gambin, Y., Lemke, E. A. & Deniz, A. A. Interplay of α-synuclein binding and conformational switching probed by single-molecule fluorescence. Proc. Natl. Acad. Sci. USA 106, 5645–5650 (2009).

43. Cremades, N. et al. Direct observation of the interconversion of normal and toxic forms of α-synuclein. Cell 149, 1048–1059 (2012).

44. Hervás, R. et al. Common features at the start of the neurodegeneration cascade. PLoS Biol. 10, e1001335 (2012).

45. Neupane, K., Solanki, A., Sosova, I., Belov, M. & Woodside, M. T. Diverse metastable structures formed by small oligomers of α-synuclein probed by force spectroscopy. PLoS One 9, e86495 (2014).

46. Petrosyan, R., Narayan, A. & Woodside, M. T. Single-molecule force spectroscopy of protein folding. J. Mol. Biol. 433, 167207 (2021).

47. Woodside, M. T. & Block, S. M. Reconstructing folding energy landscapes by single-molecule force spectroscopy. Annu. Rev. Biophys. 43, 19–39 (2014).

48. Yu, H. et al. Direct observation of multiple misfolding pathways in a single prion protein molecule. Proc. Natl. Acad. Sci. USA 109, 5283–5288 (2012).

49. Yu, H. et al. Energy landscape analysis of native folding of the prion protein yields the diffusion constant, transition path time, and rates. Proc. Natl. Acad. Sci. USA 109, 14452–14457 (2012).

50. Yu, H. et al. Protein misfolding occurs by slow diffusion across multiple barriers in a rough energy landscape. Proc. Natl. Acad. Sci. USA 112, 8308–8313 (2015).

51. Yen, C. F., Harischandra, D. S., Kanthasamy, A. & Sivasankar, S. Copper-induced structural conversion templates prion protein oligomerization and neurotoxicity. Sci. Adv. 2, e1600014 (2016).

52. Raspadori, A. et al. Evidence of orientation-dependent early states of prion protein misfolded structures from single molecule force spectroscopy. Biology 11, 1358 (2022).

53. Ganchev, D. N., Cobb, N. J., Surewicz, K. & Surewicz, W. K. Nanomechanical properties of human prion protein amyloid as probed by force spectroscopy. Biophys. J. 95, 2909–2915 (2008).

54. Gupta, A. N. et al. Pharmacological chaperone reshapes the energy landscape for folding and aggregation of the prion protein. Nat. Commun. 7, 12058 (2016).

55. Petrosyan, R., Patra, S., Rezajooei, N., Garen, C. R. & Woodside, M. T. Unfolded and intermediate states of PrP play a key role in the mechanism of action of an antiprion chaperone. Proc. Natl. Acad. Sci. USA 118, e2010213118 (2021).

56. Qing, L. L., Zhao, H. & Liu, L. L. Progress on low susceptibility mechanisms of transmissible spongiform encephalopathies. Zool. Res. 35, 436–445 (2014).

57. Watts, J. C. et al. Evidence that bank vole PrP is a universal acceptor for prions. PLoS Pathog. 10, e1003990 (2014).

58. Nicot, S. & Baron, T. Strain-specific barriers against bovine prions in hamsters. J. Virol. 85, 1906–1908 (2011).

59. Wang, M. D., Yin, H., Landick, R., Gelles, J. & Block, S. M. Stretching DNA with optical tweezers. Biophys. J. 72, 1335–1346 (1997).

60. Dudko, O. K., Hummer, G. & Szabo, A. Intrinsic rates and activation free energies from single-molecule pulling experiments. Phys. Rev. Lett. 96, 108101 (2006).

61. Hinczewski, M., Hyeon, C. & Thirumalai, D. Directly measuring single-molecule heterogeneity using force spectroscopy. Proc. Natl. Acad. Sci. USA 113, E3852– E3861 (2016).

62. Yu, H., Siewny, M. G. W., Edwards, D. T., Sanders, A. W. Perkins, T. T. Hidden dynamics in the unfolding of individual bacteriorhodopsin proteins. Science 355, 945–950 (2017).

63. Jahn, T. R. & Radford, S. E. Folding versus aggregation: polypeptide conformations on competing pathways. Arch. Biochem. Biophys. 469, 100–117 (2008).

64. Plaxco, K. W. et al. Evolutionary conservation in protein folding kinetics. J. Mol. Biol. 298, 303–312 (2000).

65. Shrestha, S. & Clark, A. C. Evolution of the folding landscape of effector caspases. Journal of Biological Chemistry 297, 101249 (2021).

66. Lim, S. A., Bolin, E. R. & Marqusee, S. Tracing a protein’s folding pathway over evolutionary time using ancestral sequence reconstruction and hydrogen exchange. Elife 7, e38369 (2018).

67. Dima, R. I. & Thirumalai, D. Exploring the propensities of helices in PrPC to form β sheet using NMR structures and sequence alignments. Biophys. J. 83, 1268–1280 (2002).

68. Ehsani, S. et al. Evidence for retrogene origins of the prion gene family. PLoS One 6, e26800 (2011).

69. Tompa, P., Tusnády, G. E., Cserző, M. & Simon, I. Prion protein: evolution caught en route. Proc. Natl. Acad. Sci. USA 98, 4431–4436 (2001).

70. Morrissey, M. P. & Shakhnovich, E. I. Evidence for the role of PrP(C) helix 1 in the hydrophilic seeding of prion aggregates. Proc. Natl. Acad. Sci. USA 96, 11293–11298 (1999).

71. Moulick, R., Das, R. & Udgaonkar, J. B. Partially unfolded forms of the prion protein populated under misfolding-promoting conditions. J. Biol. Chem. 290, 25227–25240 (2015).

72. Singh, J. & Udgaonkar, J. B. Molecular mechanism of the misfolding and oligomerization of the prion protein: current understanding and its implications. Biochemistry 54, 4431–4442 (2015).

73. Zhou, M., Ottenberg, G., Sferrazza, G. F. & Lasmézas, C. I. Highly neurotoxic monomeric α-helical prion protein. Proc. Natl. Acad. Sci. USA 109, 3113–3118 (2012).

74. Sanz-Hernández, M. et al. Mechanism of misfolding of the human prion protein revealed by a pathological mutation. Proc. Natl. Acad. Sci. USA 118, e2019631118 (2021).

75. Vázquez-Fernández, E. et al. The structural architecture of an infectious mammalian prion using electron cryomicroscopy. PLoS Pathog. 12, e1005835 (2016).

76. Kraus, A. et al. High-resolution structure and strain comparison of infectious mammalian prions. Mol. Cell 81, 4540–4551 (2021).

77. Hosszu, L. L. P. et al. Structural mobility of the human prion protein probed by backbone hydrogen exchange. Nat. Struct. Biol. 6, 740–743 (1999).

78. Gerum, C., Silvers, R., Wirmer-Bartoschek, J. & Schwalbe, H. Unfolded-state structure and dynamics influence the fibril formation of human prion protein. Angew. Chem. Intl. Ed. 48, 9452–9456 (2009).

79. Case, D. A. et al. Amber 2018. University of California, San Francisco. (2018).

